# The scope and limits of fine-grained image and category information in the ventral visual pathway

**DOI:** 10.1101/2024.08.04.606507

**Authors:** Markus W. Badwal, Johanna Bergmann, Johannes Roth, Christian F. Doeller, Martin N. Hebart

## Abstract

Humans can easily abstract incoming visual information into discrete semantic categories. Previous research employing functional MRI (fMRI) in humans has identified cortical organizing principles that allow not only for coarse-scale distinctions such as animate versus inanimate objects but also more fine-grained distinctions at the level of individual objects. This suggests that fMRI carries rather fine-grained information about individual objects. However, most previous work investigating fine-grained category representations either additionally included coarse-scale category comparisons of objects, which confounds fine-grained and coarse-scale distinctions, or only used a single exemplar of each object, which confounds visual and semantic information. To address these challenges, here we used multisession human fMRI (female and male) paired with a broad yet homogenous stimulus class of 48 terrestrial mammals, with 2 exemplars per mammal. Multivariate decoding and representational similarity analysis (RSA) revealed high image-specific reliability in low- and high-level visual regions, indicating stable representational patterns at the image level. In contrast, analyses across exemplars of the same animal yielded only small effects in the lateral occipital complex (LOC), indicating rather subtle category effects in this region. Variance partitioning with a deep neural network and shape model showed that across exemplar effects in EVC were largely explained by low-level visual appearance, while representations in LOC appeared to also contain higher category-specific information. These results suggest that representations typically measured with fMRI are dominated by image-specific visual or coarse-grained category information but indicate that commonly employed fMRI protocols may reveal subtle yet reliable distinctions between individual objects.

**Significance Statement:** While it has been suggested that functional MRI (fMRI) responses in ventral visual cortex carry fine-grained information about individual objects, much previous research has confounded fine-grained with coarse-scale category information or only used individual visual exemplars, which potentially confounds semantic and visual object information. Here we address these challenges in a multisession fMRI study where participants viewed a highly homogenous stimulus set of 48 land mammals with 2 exemplars per animal. Our results reveal a strong dominance of image-specific effects and additionally indicate subtle yet reliable category-specific effects in lateral occipital complex, underscoring the capacity of commonly employed fMRI protocols to uncover fine-grained visual information.

## Introduction

Our visual system provides us with the remarkable ability to derive from our immediate sensory experience a deeper semantic understanding of objects, allowing us to recognize them and form higher-level categories. This ability is thought to be supported by the ventral visual pathway that comprises lateral and ventral occipitotemporal cortex (LOTC; VOTC). Beyond its characteristic responses to faces, scenes, bodies, objects and text (Kanwisher, 2010a), ventral visual cortex exhibits distinct responses differentiating between animate and inanimate objects (Chao et al., 1999; Kriegeskorte et al., 2008; Konkle & Caramazza, 2013; Hebart et al., 2023). Recent work suggests that VOTC and LOTC respond to dimensions such as animacy in a continuous, or even hierarchical, manner (Connolly et al., 2016; Sha et al., 2015; Thorat et al., 2019), indicating the presence of neural representations of increasingly complex categorical information in these brain regions that may only in part be explained by low-level visual features (Jozwik et al., 2016). Thus, while there is disagreement about the degree to which these effects reflect visual or semantic features (Andrews et al., 2010, 2015; Coggan et al., 2016; Long et al., 2018; Bracci et al., 2017, 2019; Jagadeesh & Gardner, 2022), there is a general consensus that ventral visual cortex carries representations that allow humans to distinguish objects at a coarse categorical level (Clarke, 2019), which can be seen as a first step towards unraveling the neural mechanisms underlying object recognition and understanding (DiCarlo et al., 2012; Bracci & Op de Beeck, 2023).

At the more fine-grained category level of individual, basic-level objects, numerous fMRI studies have addressed the question to what degree ventral visual cortex carries information distinguishing individual objects (Haxby et al., 2001; Spiridon & Kanwisher, 2002; O’Toole et al., 2005, Eger et al., 2008; Cichy et al., 2011; Clarke & Tyler, 2014; Iordan et al., 2015). These findings have been taken as evidence that ventral visual cortex indeed represents individual basic-level object categories beyond the immediate visual representations evoked by individual exemplars (Eger et al., 2008; Cichy et al., 2011; Clarke & Tyler, 2014). These cortical object representations appear to show a hierarchical organization paralleling the hierarchical organization of object categories, with more fine-grained category information found at higher spatial resolutions and more coarse-grained information at a more broadly-distributed spatial level (Brants et al., 2011; Bracci and Op de Beeck, 2023).

Despite the evidence for both coarse and fine-grained category representations in previous fMRI studies, most studies highlighting the generalization in ventral visual cortex across stimuli have included objects that are different not only at the basic but also at a more superordinate categorical level. For instance, some work has interweaved the difference between animate and inanimate stimuli with the difference between mammals and insects (e.g. Cichy et al., 2011; Clarke & Tyler, 2014), which are hierarchically different category levels and known to be represented quite differently (Kriegeskorte et al., 2008, Connolly et al., 2012; Hebart et al., 2020). Despite most studies being interested in object-specific effects, most of them do not distinguish between fine-grained and coarse-grained category information and run comparisons including all levels of comparisons. Thus, for studies including both coarse-scale and fine-grained category comparisons, seemingly fine-grained category effects at the level of basic objects may, alternatively, be explained by the presence of coarse categorical effects, since the reported average effects of all comparisons (e.g., decoding accuracy) also includes coarse-grained effects. Adding to this challenge, much research has not been controlling for low-level or mid-level visual features (e.g. Clarke & Tyler, 2014; Singer et al., 2023). Thus, at least part of the measured response in relation to individual categories in ventral visual cortex could be driven by such basic visual features (Andrews et al., 2010, 2015; Nasr & Tootell, 2012; Bracci & Op de Beeck, 2016; Proklova et al., 2016; Long et al., 2018). Given these two potential confounds – the mixture of coarse-level and fine-grained category information and the mix of visual and semantic information – there is currently a gap in our understanding of (1) the extent to which fMRI responses in ventral visual cortex carry fine-grained categorical information and (2) the regions that allow for such distinctions.

To address these outstanding issues more specifically, here we focused on basic-level within-category distinctions in land mammals, which constitute a large homogenous class of objects (Fig 1A). This approach allowed us to control for superordinate between-category effects (e.g. animacy) or other known, coarse categorical distinctions (e.g. mammals, insects, birds, or fish). By focusing only on land mammals with faces, we implicitly controlled also for the face-body distinction that could in part be driving overall responses (Kriegeskorte et al., 2008; Ritchie et al., 2021; Proklova & Goodale, 2022). To control for basic visual features, we used two exemplars per object category, which allowed us to compare brain responses for individual images with responses across images of the same category. Finally, to maximize statistical power, we conducted our study across three scanning sessions.

**Figure 1:**
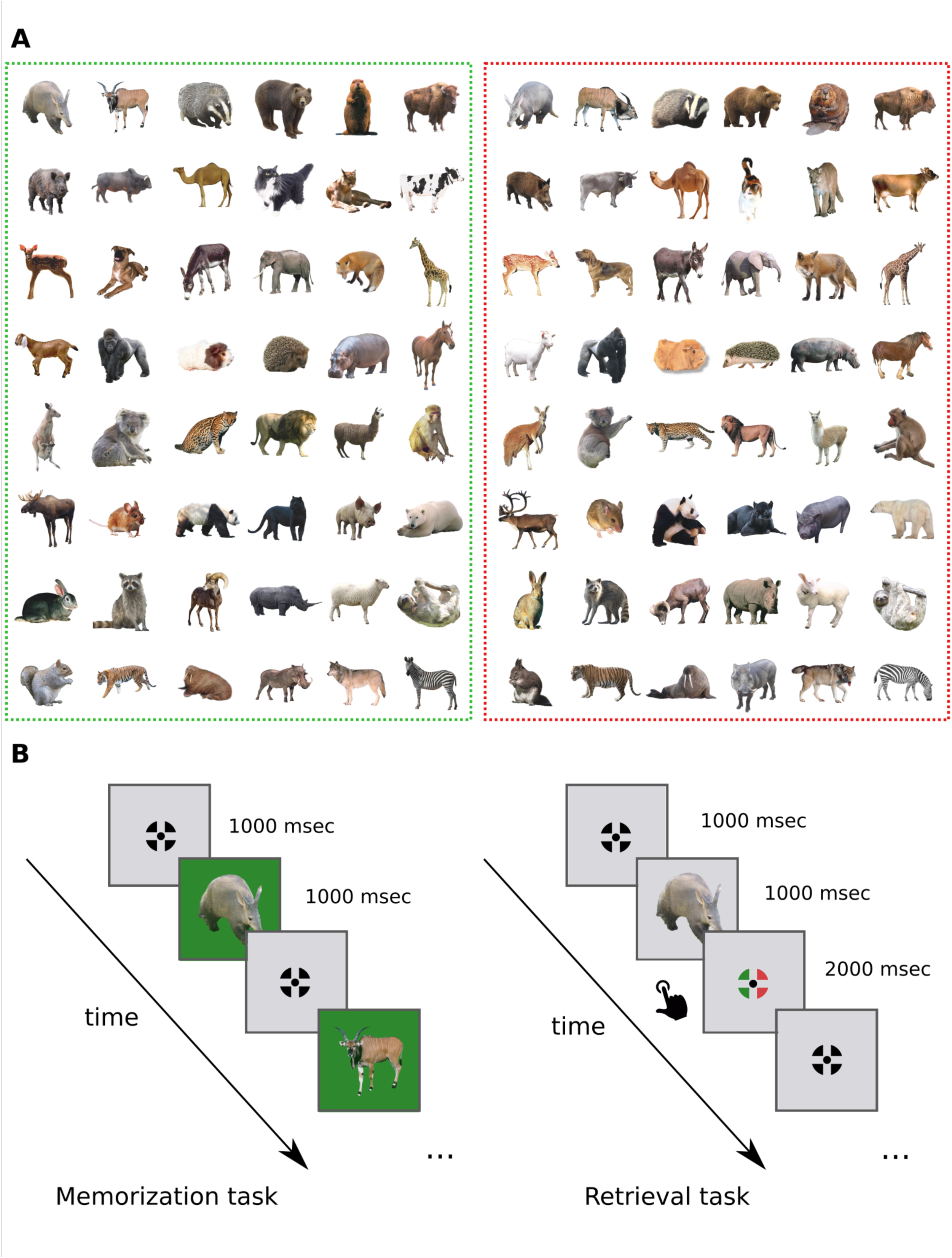
Stimulus set and experimental paradigm. ***A)*** Stimulus set: The 96 terrestrial mammal images used in the current study. Each of the 48 animals is represented by two different exemplars. Each exemplar was assigned to one of two color contexts, either green (context 1, left) or red (context 2, right), indicated by the colored rectangle. Images were collected via web search with public domain licenses. Animals are shown in alphabetical order. **B)** Experimental paradigm: The experiment consisted of a memorization task (left) and a retrieval task (right). In the memorization part, participants passively viewed the stimuli in their respective color context and were asked to memorize each exemplar. In the retrieval part, participants viewed the stimuli on a gray background. Following each presentation, the participants were asked to assign the previously seen stimulus to the correct color context with a button press.

## Materials and Methods

### Participants

For our fMRI experiment, we collected data from a total of 21 adult volunteers. Of these, one volunteer was excluded due to strong head motion and five volunteers due to poor behavioral performance (see below). Thus, our final results are based on the behavioral and fMRI data of 15 volunteers (N = 15, 7 female, 8 male, mean age: 27.8 years, age range: 22 - 38 years). The final sample size was limited by the extensive resources required for the acquisition of multisession fMRI and the limited availability of scanning slots.

All volunteers had normal or corrected-to-normal vision including normal color vision and provided written informed consent. The study was approved by the local Ethics Committee of the University of Leipzig, Germany (481/19-ek). Participants were remunerated for the online training and MRI scanning sessions.

### Stimulus set

A total of 96 object images were used in the current study, where all images were from the animal subcategory of terrestrial, non-flying mammals, i.e. mammals that live on the land (Fig. 1A). The images covered a total of 48 mammals, where each mammal was presented as two different exemplars. Each of the two exemplars had previously been assigned to one of two color contexts (green or red background), resulting in 48 mammal images in the green color context (= context 1) and 48 mammal images in the red color context (= context 2). The assignment of exemplars to color context was consistent across participants. Mammal images were collected via web search with public domain licenses. Given the comparably small number of freely available public domain images at the time of data collection, we did not employ specific criteria for finding images that were visually particularly dissimilar. Images were next edited to remove the background. For simplicity, in the following we may refer to terrestrial, non-flying mammals as animals.

As has been shown previously, models that aim to explain distinct neural representations of animate objects appear to be driven significantly by effects such as face-body distinctions, degree of animacy, and taxonomy (Connolly et al., 2012, 2016; Ritchie et al., 2021; Proklova & Goodale, 2022). In order to avoid these effects and to create a stimulus set with a high degree of homogeneity, we only included animals of the biologically distinct class of mammals. Moreover, we also excluded purely aquatic mammals or flying mammals. Thus, all animals were predominantly terrestrial, belonged to the same taxonomic class of mammals, and all images showed both faces and bodies of animals. Thus, rather than constructing a maximally diverse dataset (e.g. Hebart et al., 2020; Proklova & Goodale, 2022; Jozwik et al., 2022), we chose to control for multiple factors and created a large, highly homogenous stimulus set. This was a deliberate decision, as we were interested in the disentanglement of fine-grained representations in the ventral visual pathway.

Our choice of terrestrial mammals was driven in part by the results from Hebart et al. (2020), who showed that, based on behavioral similarity data, mammals strongly cluster together within the kingdom of animals (Hebart et al., 2020), and in part by evidence showing that highly animate objects (e.g. mammals) elicit strong and distinct neural responses (Sha et al., 2015; Proklova & Goodale, 2022).

In the MRI scanner, participants were shown the colored objects in the center of a uniform mid-gray screen through a mirror attached to the head coil, while the images were projected centrally measuring 12° × 12° visual angle onto a rear-projection screen located at the back of the scanner.

### Experimental design

The experiment consisted of two parts, a memorization and a retrieval task, performed on four subsequent days. On day 1, participants learned the stimuli and the experimental design in a one-hour behavioral online training performed at home. On days 2 to 4, participants performed a short 15 min pre-scan behavioral training and subsequently entered the scanner for data collection.

The color context memory component was included in the task as one motivation of the study was to maximize the chance of attaining relevant signal not only in posterior ventral visual cortex but also in entorhinal and hippocampal cortex, which are known to be involved in context retrieval (Libby et al., 2019; Julian & Doeller, 2021). Additionally, the design made the task sufficiently challenging to be performed over multiple days and kept participants engaged.

The online and pre-scan behavioral training consisted of the memorization task and the retrieval task (Fig. 1B) performed in alternation. During the memorization task (Fig. 1B, left), stimuli within their color context were presented at rapid succession (stimulus duration: 1000 ms; interstimulus interval: 1000 ms) while subjects were asked to memorize each image exemplar in its respective context. This part of the experiment was a passive viewing task. Each of the 96 stimuli was presented once per run. During the retrieval task (Fig. 1B, right), stimuli were again presented in fast succession (stimulus duration: 1000 ms, interstimulus interval: 3000 ms), but now displayed on a uniform gray background. During the image presentation, a black fixation crosshair (Thaler et al., 2013) appeared at the center of the screen. Within the interstimulus interval, subjects had the first 2000 ms to assign the previously displayed animal stimulus to its correct color context by clicking the right or left button depending on the response mapping indicated by the color of the fixation crosshair (Fig. 1B, right). For example, when the fixation crosshair was green on the left and red was the correct response, participants were supposed to push the right button. The side of each color on the fixation cross was randomized. The purpose of the response mapping was to decorrelate choices and button presses. In the online and pre-scan training, memorization and retrieval tasks alternated. The online training consisted of eight runs, the pre-scan training of two runs.

During the three fMRI scanning sessions, participants performed only the retrieval task, i.e. without a colored background (Fig. 1B, right). For improved motivation, participants received feedback after each run on the percentage of correct trials. Each fMRI session consisted of eight experimental runs, where each run consisted of 96 trials. Each stimulus was presented once per run. In order to improve model fitting, the event-related sequence was optimized across the three scanning sessions so that the order was maximally different between all 24 runs. All in all, there were a total of 96 trials per run, all of which were target trials, eight runs per session, and three sessions over three consecutive days, resulting in 2,304 trials for each participant and 24 repeats of each individual stimulus.

For both the memorization and the retrieval task, stimulus presentation and control were performed via a PC computer running PsychoPy2 (Peirce et al., 2019), built upon the open-source programming language Python. The online behavioral training was performed using the *Pavlovia* website (https://pavlovia.org/) build upon PsychoPy.

To minimize the chance that results were adversely affected by behavioral performance, we decided to only include participants in our analyses that reached a certain behavioral threshold. On day one, participants needed to reach at least 80% correct responses in a majority of the runs. On day two, we set the threshold at 90% and above. All participants who reached those thresholds on the first two scanning days achieved above 95% correct trials on day 3, while most were able to get at least one perfect run with 100% correct (see Results).

### MRI data acquisition

All MRI data were acquired using a 32-channel head coil on a research-dedicated Siemens Magnetom Prisma 3T MRI scanner (Siemens, Erlangen, Germany) located at the Max-Planck-Institute for Human Cognitive and Brain Sciences in Leipzig, Germany. The scanning procedure was the same for all three scanning days, with the exception of an anatomical T1-weighted scan on the first day. For the functional scans, whole-brain images were acquired using a segmented k-space and steady-state T2*-weighted multi-band (MB) echo-planar imaging (EPI) single-echo gradient sequence that is sensitive to the BOLD contrast (60 slices in interleaved ascending order; anterior-to-posterior (A–P) phase-encoding direction; TR = 1500 ms; 283 TRs per run; echo time (TE) = 24.2 ms; voxel size = 2 × 2 × 2 mm; matrix size = 96 × 96; field of view (FOV) = 204 × 204 mm; flip angle (FA) = 80°; MB acceleration factor 2, GRAPPA factor 2, partial Fourier 6/8). Each MRI session included eight functional task runs. Each functional task run was 7 min 4.5 s in length, during which 283 functional volumes were acquired. To perform distortion correction, for each functional run we recorded phase-encoding-reversed pairs. These images were incorporated in the preprocessing flow. At the end of the first scanning session, high-resolution T1-weighted anatomical sequences (MP2RAGE) were obtained from each participant to allow co-registration and brain surface reconstruction (176 slices; TR = 2300 ms; echo time (TE) = 2.98 ms; flip angle (FA) = 9 degrees; inversion time (TI) = 900 ms; matrix size = 192 × 256; FOV = 192 × 256 mm; voxel size = 1 × 1 × 1 mm).

### MRI data preparation and preprocessing

We arranged our data according to the Brain Imaging Data Structure (BIDS; Gorgolewski et al., 2016) specification in order to allow for reproducibility of the task-based fMRI study, to comply with standardization, and to use *fMRIPrep* for preprocessing.

Results included in this manuscript come from preprocessing performed using *fMRIPrep* version 21.2.0 (Esteban et al., 2019), a *Nipype* (Gorgolewski et al., 2011; Gorgolewski et al., 2017) based tool. Many internal operations of *fMRIPrep* use *Nilearn* (Abraham et al., 2014), principally within the BOLD-processing workflow. For more details of the general *fMRIPrep* pipeline and the detailed steps performed by *fMRIPrep* 21.2.0 on the anatomical and functional data of each participant see https://fmriprep.org/en/stable/workflows.html.

Functional images were not smoothed for any analysis performed in this study.

### GLM fitting

The blood-oxygen-level-dependent (BOLD) signal for each stimulus for each trial, at each voxel, was modeled separately using the least squares separate (LS-S; Mumford et al., 2012) approach. Using this approach, the general linear model (GLM) includes a single regressor for the stimulus to be estimated while all other stimuli are combined into a single nuisance regressor. Thus, for each run there are as many separate linear models fitted as there are trials (96 GLMs per run). We used LS-S on the grounds that our experiment was a rapid event-related task design with strongly overlapping evoked BOLD signals of adjacent trials, which would result in highly autocorrelated beta estimates, as well as the fact that we presented each stimulus only once per run. Mumford et al. (2012) showed that LS-S model fitting is an improved approach in comparison to a regular GLM for obtaining reliable activation patterns for rapid event-related designs like ours. Each onset regressor was convolved with a canonical hemodynamic response function. For the experimental runs, we additionally included six head movement regressors derived from estimated head motion parameters (translation and rotation along the x-, y-, and z-axes) in our GLMs. After fitting the GLM, our analysis yielded voxelwise beta weights for each stimulus for each run. For all analyses reported below, the estimated beta weights for each stimulus within each session were averaged across all 8 runs, resulting in a single beta weight for each stimulus for each session in each voxel. Our rationale behind averaging the beta weights within sessions was that every stimulus was presented only once per run, leading to noisy fitting. Thus, the averaged session-wise beta weight provided a more stable and reliable estimate. Thus, we ended up with a total of 96 beta weights for each of the three scanning sessions, one for each animal stimulus, for each subject. These voxelwise beta weights were used as input for all subsequent voxel pattern analyses and model comparisons.

fMRI data model fitting was done using custom code in Python partly based on the *Nilearn* toolbox (https://github.com/nilearn/nilearn). We performed model fitting in native space as many of our analyses were carried out within ROIs in each individual participant.

### ROI definition

To quantify BOLD effects at the regional level, we defined regions of the ventral visual pathway by using ROIs from the probability brain maps of Wang et al. (2015), selected masks from n*eurosynth* (https://neurosynth.org/) as well as *neurovault* (https://neurovault.org/), and selected masks available in *FSL* (Fig. 2). We included Hippocampus (HC) and perirhinal cortex (PRC) as an extension of the ventral visual pathway, as recent studies have shown that they are implicated in higher conceptual representations (Clarke & Tyler, 2014; Theves et al., 2020). All ROIs are bilateral clusters including the left and right hemisphere.

**Figure 2:**
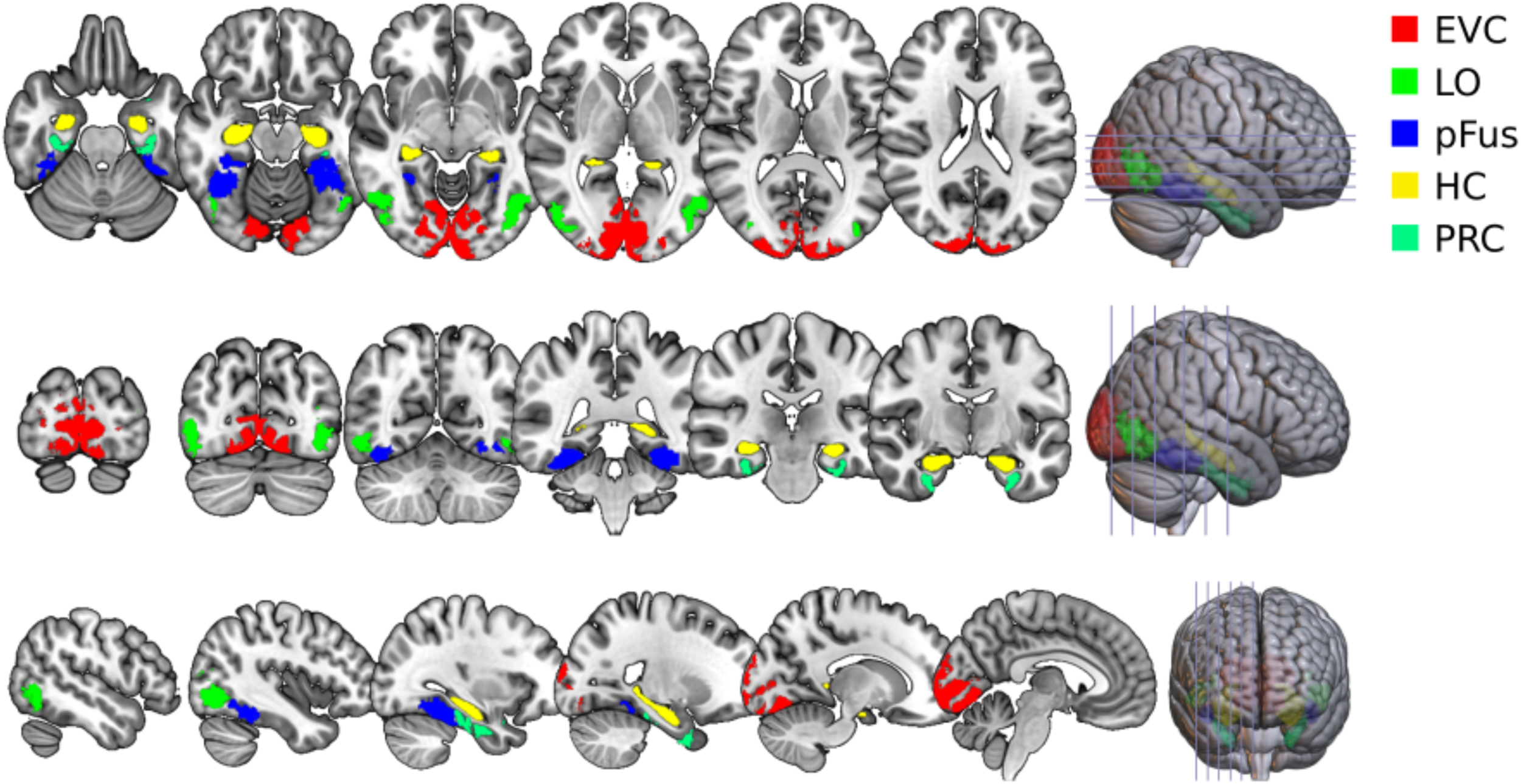
Definition of the ROIs. We defined the following ROIs to describe the ventral visual pathway including extended ROIs: EVC (V1 – V3), lateral occipital cortex (LO), posterior fusiform (pFus), hippocampal cortex (HC) and perirhinal cortex (PRC). LO and pFus are together referred to as lateral occipital complex (LOC).

We merged V1, V2 and V3 from Wang et al. (2015) to construct an early visual cortex (EVC) mask (27,805 mm^3^). Splitting the lateral occipital complex (LOC), in two subregions, the lateral occipital region (LO; 9,263 mm^3^) as well as the posterior fusiform region (pFus; 11,273 mm^3^) were constructed using the object association regions from *neurosynth* and selecting the appropriate subregions. The HC mask (8,719 mm^3^) was taken from the Harvard-Oxford brain atlas included in *FSL,* and the PRC mask (7,910 mm^3^) from a probabilistic map constructed of 38 healthy subjects (https://neurovault.org/images/60340/).

For all ROIs, we set a threshold of ≥50 % of the maximum original mask values to only include mostly unambiguous voxels. For analyses, all ROIs were shifted from MNI152 space into the native space of each participant using the transformation matrix of the T1-weighted image provided by *fMRIPrep* and resampled to the nominal voxel size of the functional volumes.

### Reliability-based voxel selection

In order to maximize sensitivity of our analyses across sessions, we used reliability-based voxel selection (Tarhan & Konkle, 2020), which selects a subset of voxels within each ROI that exhibit a stable voxel response profile across conditions. Accordingly, an array containing the beta estimates of all stimuli was computed for each voxel for each session (96 beta estimates per session in the current study). The Pearson correlation between these arrays was then computed for each voxel across all pairs of the three sessions, and the mean correlation taken as the overall reliability value. Considering that we computed the reliability over three sessions, as opposed to two, we used the Spearman-Brown prediction formula (Spearman, 1910; Brown, 1910) with a *k*-Factor of 1.5 to correct for the reliability estimates across all three sessions (see below).

We ended up including all voxels in further analyses of which the estimated correlation exceeded a threshold of *r* ≥ 0.15. Based on visual inspection, this threshold optimized the natural tradeoff between brain coverage and stability of the data. Please note, though, that the results were highly stable irrespective of this voxel selection strategy. Using reliability-based voxel selection allowed us to select the most informative and stable voxels across all three sessions in the multi-voxel pattern analyses (MVPA; Haxby et al., 2001) while discarding noisy voxels, thus maximizing the chance of identifying reliable effects across sessions and conditions.

### Multivariate decoding analysis

We used multivariate decoding analysis on the fMRI multivoxel patterns (Haxby et al., 2001; Kriegeskorte et al., 2006; Haynes & Rees, 2005) to determine to what extent image-specific and category-specific information was encoded along the ventral visual pathway. A linear support vector machine (SVM) classifier was trained either with a focus on image-specific representations (training and testing within one image set) or on category-specific representations (training on one image set and testing on the other, and vice versa). All analyses were conducted independently for each subject and each ROI. For a given pair of categories, multivoxel pattern vectors from 2 out of 3 sessions were assigned to the training data, and the left out session was used as the independent test data. Data were cross-validated using 6-fold cross-validation, comprising all possible combinations of training and testing sessions. This analysis was repeated for all pairs of objects, and results were averaged across all comparisons. Group-level analyses of decoding accuracy across all subjects was conducted by means of paired t-tests or one-sample t-tests relative to the classification chance level of 50 %.

### Representational similarity analysis: Reliability of the representational geometry across sessions and exemplars and model comparisons

Representational similarity analysis (RSA) was used (1) to estimate the stability of the pairwise neural activity patterns across sessions, (2) to compare the neural representations across exemplars of the same concept, and (3) to estimate the representational similarity between neural activity and various layers of different deep neural networks as well as semantic embedding models. All RSA was conducted using custom code in Python largely based on the open-source toolbox *rsatoolbox* (rsatoolbox.readthedocs.io; Schütt et al., 2023).

Neural representational dissimilarity matrices (RDMs) for each subject for the different ROIs as well as model RDMs were created using 1 minus Pearson’s correlation (*r*) as the similarity metric between the representational patterns of the stimuli. We used 1 minus Pearson correlation as the distance metric since it is commonly used and also applied in previous studies of animacy (e.g. Connolly et al., 2016; Bracci et al., 2019). In addition, Pearson’s r showed higher stability in across session representations than Euclidean or cosine distance (Kriegeskorte et al., 2008b). We calculated the neural RDMs by calculating the pairwise correlation distance of stimulus-specific beta patterns, yielding a 96 × 96 fMRI RDM for each ROI for each participant. For the various DNNs (see below), we constructed layer-specific RDMs based on 1 minus the Pearson’s correlation between all pairs of vectors of unit responses to each stimulus. For each semantic embedding model (see below), a single RDM was constructed based on the word embedding of each concept. *T*o compare RDMs from different sessions and across modalities, we took the lower triangular part of each matrix excluding the diagonal (Ritchie et al., 2017) and converted them to vectors. Subsequently, these vectors were compared using Spearman’s rank correlation rho for pairwise RDM comparisons.

Since we acquired functional data from three separate sessions across three subsequent days, we sought to determine how reliable the within-exemplar representational relationships were across sessions. To quantify the reliability, we computed Spearman’s rho between the context-specific RDMs containing the respective 48 stimuli (48 × 48 RDM) of each ROI across all three sessions. The correlation values were then averaged across all comparisons. This way, we obtained a single correlation value for each subject for each ROI that indicated the reliability of the representational geometry across sessions. We used the context-specific RDMs for within-exemplar correlations in order to have the same modality (48 × 48 RDM) as when looking at across-exemplar similarities.

Having two exemplars of each concept in our data set, we further wanted to test whether there are generalizable representational relationships between different exemplars of the same animal species (e.g. Is the representational relationship between a dog and a mouse generalizable, independent of its visual appearance?). We thus computed the Spearman’s rho between the two context RDMs (48 × 48 RDM), correlating context 1 RDM (green) with context 2 RDM (red) across all three sessions and took the average. This way, we obtained a single correlation value for each subject for each ROI that indicated the representational similarity between the two exemplars of all concepts.

### Variance Partitioning: Modeling visual, semantic and task effects

To determine the role of visual features, semantics, and task in the estimated representational similarities, we carried out variance partitioning to partial out variance explained by these factors (e.g. Hebart et al., 2018; Cichy et al., 2019). Once an effect has been partialled out, a remaining significant correlation between the residual RDMs would indicate that the partialled-out factors cannot fully explain the hypothesized relationship between the RDMs.

Deep neural networks (DNNs) reflect an established class of visual models of neural representations along the ventral visual pathway (Khaligh-Razavi & Kriegeskorte, 2014; Cichy et al., 2016) as well as perceived similarity between images (Kubilius et al., 2016, Hebart et al., 2022). We chose AlexNet (Krizhevsky et al., 2012) as it has been used commonly in the field of cognitive computational neuroscience to model the various processing stage of object recognition (e.g. Cichy et al., 2016; Bracci et al., 2019; Cichy et al., 2019; Mehrer et al., 2021). The THINGSvision toolbox (Muttenthaler & Hebart, 2021) was used to extract layer-specific activity patterns from AlexNet, which had been pretrained on the ImageNet dataset (Deng et al., 2009). Based on previous findings (e.g. Cichy et al., 2016; Bankson et al., 2018), we modeled low-level visual features using an early convolutional layer (= layer #2). We calculated the pairwise Pearson’s *r* between layer-specific vectors, generating three distinct RDMs: a 96 × 96 RDM for the entire stimulus set, a 48 × 48 RDM for the stimuli in context 1, and a 48 × 48 RDM for the stimuli context 2. In addition, to model mid-level visual features, we used the “ShapeComp” model to account for such visual feature effects, which is a model of perceived shape similarity (Morgenstern et al., 2021). Here again, we calculated the pairwise Pearson’s r between shape feature vectors, generating a distinct 96 x 96 RDM indicating the pairwise shape similarity between all stimuli.

To assess fine-grained information about the semantic meaning of the shown stimuli, two similarity models were used. As a first model, we included the THINGS embedding of perceived similarity of Hebart and colleagues (2020), which had been trained on behavioral data during a triplet odd-one-out task (Hebart et al., 2020, 2023) and which captures behaviorally-relevant visual and conceptual features. Here, we generated the similarity matrix from the embedding, assuming the similarity as the probability of not choosing a pair together across all possible pairs of comparisons, and focused the possible triplets in the comparison on the set of 48 objects (48 × 48 RDM). As a second model, we used the CSLB concept property norm model (Devereux et al., 2014) to include sets of semantic features explicitly listed by human participants. In this model, the semantic features are labeled to belong to a certain semantic feature type (e.g. external component, behavior), which allows for the subselection of features. Each object vector consists of binary numbers that reflect the presence or absence of a semantic feature. Since the CSLB model contains only 34 of the 48 basic-level concepts in our study, we limited our analyses to these concepts when using the CSLB semantic feature model for comparison. We extracted the semantic feature vectors for each concept and calculated the Pearson’s r distance between each pair of vectors (34 × 34 RDM).

### Statistical analysis

For testing effects within each ROI, we used a two-tailed one sample t-test on the group results with an expected population mean of zero (e.g. testing whether the activity patterns are reliable in a ROI). For testing effects between ROIs, we used a pairwise t-test (e.g. difference between reliability in two ROIs). Where necessary, we performed a non-parametric permutation test and indicated this in the text. All reported p-values were corrected for multiple comparisons across ROIs using Bonferroni correction. Results are reported as mean values with the respective standard error of the mean (SEM, ±).

## Results

Our goal in this study was to determine the degree to which members of a highly homogenous taxonomic subgroup, in our case terrestrial mammals, would elicit distinct voxel activation patterns. More specifically, we aimed to (1) identify the degree to which differences between basic-level objects could be identified that would generalize across different exemplars of the same mammal, and (2) to identify whether a detectable semantic relationship within such a homogenous stimulus set could be found.

### Behavioral results

The strict inclusion criteria used in the present study (see Material & Methods) ensured (1) that random guessing was unlikely and (2) that volunteers had a reliable grasp of each stimulus in its respective color context and were sufficiently attentive. This is reflected in the behavioral results. Participants performed very well on all three days, with an accuracy that reached very high levels on the second day (Day 1: 85.02% ± 6.82, Day 2: 95.63% ± 3.80, Day 3: 98.13% ± 2.34). We also found a near absence of missed responses in the 2000 ms interstimulus response window across all three scanning days (Misses: 3.15 ± 2.98 out of 2,304 trials), corroborating the evidence that participants were paying attention and were alert throughout.

### Reliability of voxel response profiles decreases along the ventral visual pathway

Having verified that participants had a reliable apprehension of the stimuli, we were next interested in the degree to which each participant’s fMRI data exhibited reliable activity patterns across the three different scanning days. To test this, we used reliability-based voxel selection (Tarhan & Konkle, 2020), while at the same time maximizing statistical power across sessions and discarding noisy voxels from comparisons. A qualitative illustration of the results shows that early visual regions contained many highly reliable voxels, with a decline along the ventral visual pathway (Fig. 3a), which is in line with previous findings (Tarhan & Konkle, 2020). A quantitative depiction confirms that EVC contains the largest fraction of reliable voxels out of all ROIs (EVC reliability: 49.73% ± 2.29%, 1367 voxels ± 63; Fig. 3b). In the lateral occipital complex, we also found large fractions of voxels that survived our inclusion threshold (LO reliability: 37.27% ± 2.25%, 316 voxels ± 19; pFus reliability: 15.63% ± 1.34%, 174 voxels ± 15; Fig. 3b). Since the fraction of reliable voxels and reliability measures decreased significantly as compared with EVC, it is reasonable to assume that the information encoded in LO and pFus is more difficult to capture than in early visual regions, possibly also due to the reduced MRI signal in these regions. Inspecting the brain regions considered to be at the apex of the ventral visual pathway, we found only a small number of voxels that showed reliable response profiles in HC and PRC (HC reliability: 6.13% ± 0.54, 52 voxels ± 5; PRC reliability: 5.85% ± 0.42, 20 voxels ± 2; Fig. 3b). This could at least in part be explained by a combination of signal loss and distortions, which are common in these regions (Weiskopf et al., 2006; Olman et al., 2009). Additionally, the information encoded by the neurons in these regions might be too complex to be captured by voxel patterns recorded with fMRI.

**Figure 3:**
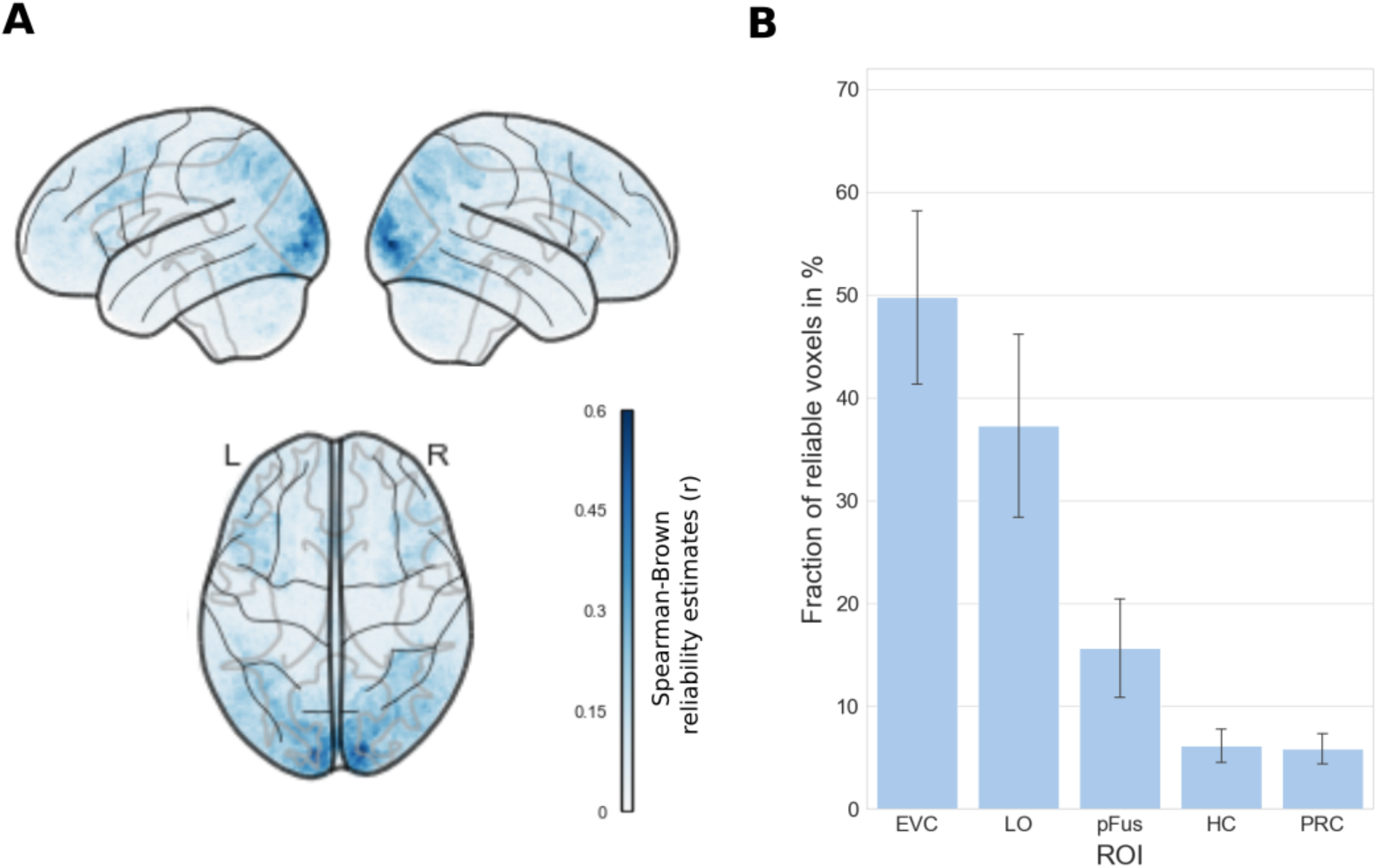
Voxel reliability. **A)** Glass brain display of the whole-brain Spearman-Brown reliability estimates (unthresholded) averaged across all subjects using the reliability-based voxel selection method (Tarhan and Konkle, 2020). **B)** Barplot display of the fraction of reliable voxels exceeding our threshold of r ≥ 0.15 in percentage (y-axis) in each ROI (x-axis). Error bars reflect 95% confidence intervals.

In summary, there is large coverage of reliable voxels in the regions along the ventral visual pathway, with many highly reliable voxels in visual areas and a significant decrease of reliability towards the more anterior-ventral regions (Fig. 3a, b).

### Decoding of image-specific and category-specific information

Our primary aim was to reveal the degree to which ventral visual cortex contained representations within the highly homogenous stimulus class of mammals that would generalize beyond image-specific effects. To this end, we trained and tested a linear classifier on image-specific and category-specific multivoxel patterns, respectively. We operationalized category-specific effects as information that would generalize across visually distinct exemplars, resulting in above chance cross-decoding accuracy for visually dissimilar exemplars.

The image-specific and category-specific decoding results are shown in Figure 4a. We found that image-specific information could be decoded significantly above chance in all ROIs along the ventral visual pathway. Especially in early and high-level visual regions, image-specific decoding accuracies were very high (EVC: 89.54% ± 1.05%, LO: 76.52% ± 0.98%, pFus: 76.69% ± 0.70%, p < 0.001). This result was expected as these regions are known to represent low- and high-level visual features of the viewed images. Further, voxel patterns in more anterior brain regions still contained sufficient image-specific information to accurately decode single images significantly above chance level (HC: 71.18% ± 0.77%, p < 0.001; PRC: 67.68% ± 0.60%, p < 0.001). The observed phenomenon likely suggests that the strong visual features effects from visual regions extended to higher brain regions along the ventral visual pathway, exerting a predominant influence on the representational patterns.

**Figure 4:**
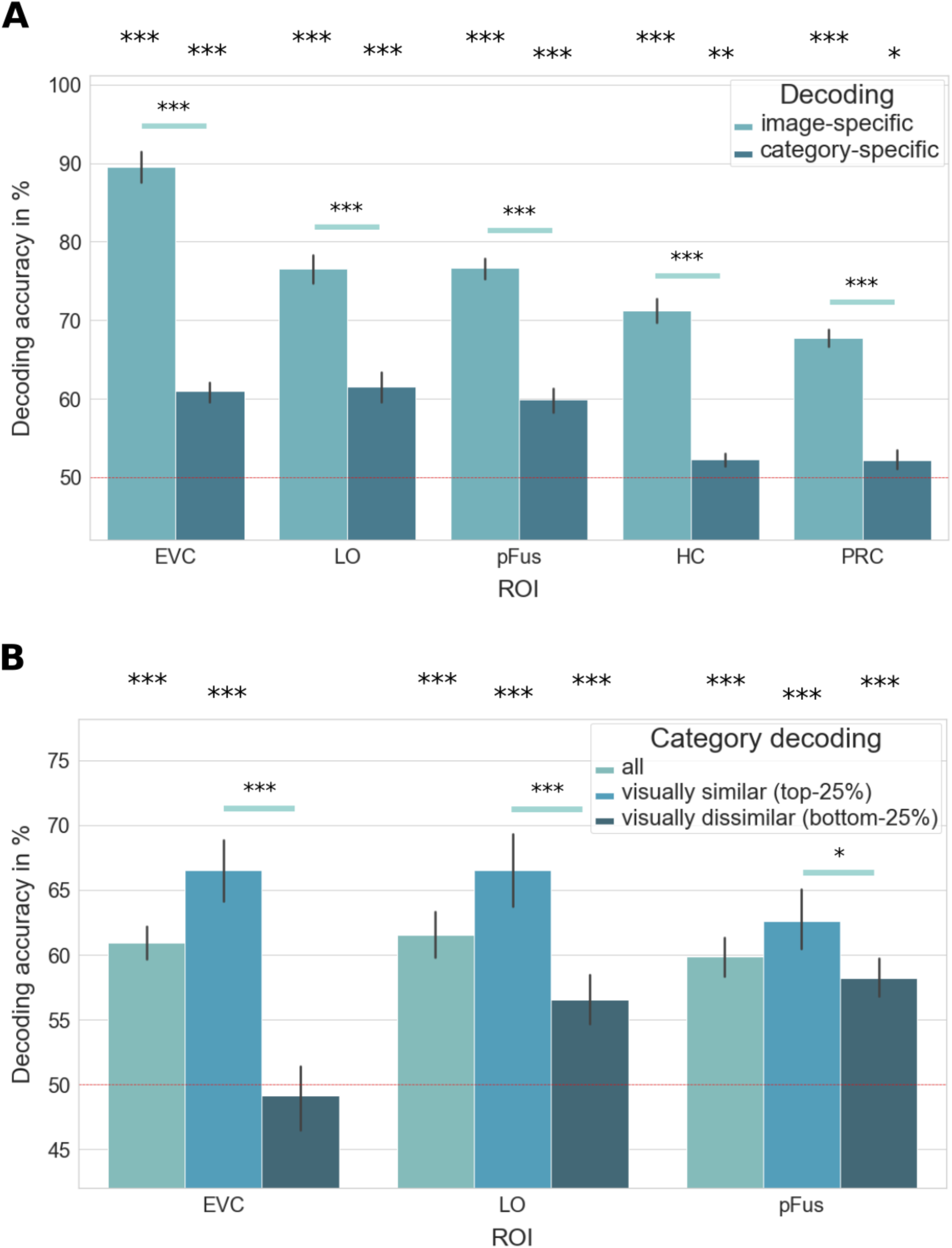
Decoding accuracy for an image-specific and category-specific SVM. **A**) ROI-wise decoding accuracies of pairwise comparisons at the image-specific level (training and testing within exemplar) and the category-specific level (training and testing across exemplars of the same object). The red line indicates chance-level for pair-wise decoding (50%). **B**) Significant ROI-wise category-specific effects were additionally split for visual similarity, as defined by the representational similarity within an early convolutional DNN layer. Results reflect the top quartile and bottom quartile of visual similarity. Asterisks indicate significance when compared with zero at significance levels: *** p < 0.001, ** p < 0.01, * p < 0.05, Bonferroni-corrected. Error bars reflect 95% confidence intervals.

To identify the degree to which higher category-specific information is present in each ROI, we implemented cross-decoding (Fig. 4a). The reasoning for adopting this approach is that, if the classifier trained on one exemplar is able to accurately decode another exemplar from the same basic level category, it can be inferred that the fundamental representational structure of the two visually dissimilar but categorically similar objects is significantly related. For cross-decoding, we found above chance accuracy in EVC (60.95% ± 0.66%, p < 0.001), LO (61.53% ± 1.00%, p < 0.001) and pFus (59.85% ± 0.88%, p < 0.001). Decoding accuracy in HC (52.25% ± 0.44%, p = 0.001) and PRC (52.20% ± 0.63%, p = 0.019) exceeded chance level, but with comparably small effects. Due to the numerically small effects in HC and PRC, we did not carry out follow-up analyses on these regions.

Above chance decoding of visually different exemplars of the same basic-level category could hint at higher level categorical information encoded in the voxel patterns. Alternatively, it is possible that exemplars of the same basic-level category still share distinct visual features that make them more similar to each other than to exemplars from other categories allowing for above chance decoding. This idea is supported by the above-chance decoding accuracies in EVC. To test this alternative explanation, we split the categories in different subgroups based on their visual similarity. Visual similarity was computed as a proxy, using Pearson similarity of the first convolutional layer of AlexNet (see Material & Methods). We then focused our analyses on the top- and bottom-quartile of stimuli according to their visual similarity. We then repeated the category-specific decoding analysis but focused all comparisons on object pairs within these quartiles. The results are displayed in Figure 4b. The decoding accuracy was higher for the visually most similar exemplars than for the visually most dissimilar exemplars in all ROIs. This difference is highly significant in EVC and LO (EVC: mean difference = 17.42%; LO: mean difference = 10.00%, p < 0.001) and significant in pFus (mean difference = 4.39%, p = 0.015), suggesting that visual similarity is a central factor in all ROIs. In EVC, decoding accuracy for the visually most dissimilar categories dropped to chance level (49.12% ± 1.24%), indicating that the visual similarity comparison effectively captured visual effects. In LO and pFus, decoding accuracy for the visually most dissimilar categories remained significantly above chance (LO: 56.54% ± 0.95%; pFus: 58.16% ± 0.78%) indicating category information beyond the basic visual effects captured by the early DNN layer.

Taken together these results show that visually identical images can be reliably decoded from all ROIs along the ventral visual pathway and that visual similarity dominates the decoding accuracy between different exemplars of the same basic-level category.

### Reliability of representational geometry within and across animal exemplars

Next, we used RSA to compute the across-session reliability of the representational geometry within and across exemplars. While multivariate decoding revealed across exemplar category-specific information contained in the multivoxel patterns, RSA allows to reveal whether the global representational geometry is similar between two sets of animal exemplars, which would more broadly indicate generalizable representations.

As participants were exposed to visually identical stimuli during each scan, we predicted the elicited fMRI patterns within-exemplars in early- and mid-level visual areas to be relatively similar across sessions. Indeed, the reliability in EVC was highly significant when compared with zero (mean *r* = 0.38 ± 0.01, *p* < 0.001, Fig. 5b) and significantly higher than in any other ROI (*p* < 0.001). We were further interested in the extent to which the representations in higher visual areas in occipito-temporal cortex (OTC) showed reliable activity patterns. These representations were expected to encode more complex, possibly semantic, features of object identity (e.g. Konkle & Oliva, 2012; Bracci & Op de Beeck, 2016; Contier et al., 2023). Indeed, the reliability in LO as well as pFus was significantly larger than zero (LO: mean *r* = 0.19 ± 0.01, pFus: mean *r* = 0.19 ± 0.01, *p* < 0.001, Fig. 5b), with no significant difference between the two (*p* = 0.81). This indicates that the information contained in the representational geometry in LOC can be reliably captured across multiple sessions. Finally, we examined the pattern reliability in the final and most anterior regions of the visual processing hierarchy, which are thought to code for concept-specific information (Clarke & Tyler, 2014; Theves et al. 2020). Even though low, the across-session reliability in these regions remained significant when compared to zero (HC: mean *r* = 0.02 ± 0.004, PRC: mean *r* = 0.05 ± 0.008, *p* < 0.001; Fig. 5b). Together, these results suggest that the further one moves along the hierarchy of the ventral visual stream, the lower the reliability of the representational geometry between objects within a highly homogenous stimulus set. This decline may be explained by a combination of increasingly more complex information encoded in the representational patterns and a decrease of fMRI signal quality. These results are in line with our analyses of the voxel reliability measures (Fig. 3).

**Figure 5:**
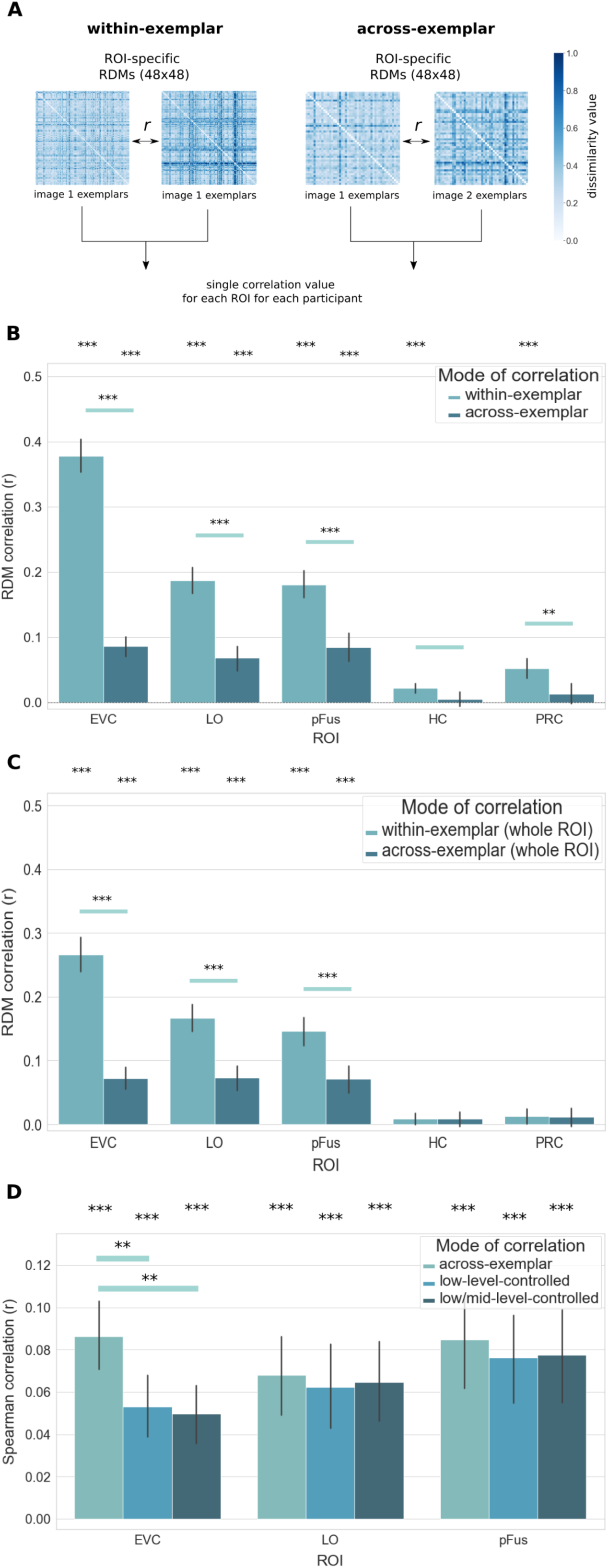
Reliability of representational similarity within and across exemplars. **A)** Analysis approach for comparing representational similarity within and across exemplars, with reliability estimated across scanning sessions. **B)** Pattern reliability within and across exemplars, estimated across scanning sessions, where across exemplar comparisons reflect a representational geometry that is abstracted away from individual images. **C)** Pattern reliability within and across exemplars but without reliability-based voxel selection, showing overall smaller effects yet a similar pattern of results. This suggests that reliability-based voxel selection did not induce any specific bias. **D)** Comparison of the across-exemplar correlation effects before and after solely partialling out the early convolutional layer RDM from AlexNet (“low-level-controlled”) as well as after additionally partialling out the shape model (“low/mid-level-controlled”). Results are shown only for ROIs with significant across exemplar reliability. Asterisks indicate significance when compared with zero at significance levels: *** p < 0.001, ** p < 0.01, * p < 0.05, Bonferroni-corrected. Error bars reflect 95% confidence intervals.

Having established the within-exemplar reliability as a noise ceiling, next we investigated whether the representational similarity generalizes across exemplars, which would hint at a basic category-specific representational geometry. To identify such categorical representations, we computed separate RDMs for each of the two contexts for each session within each participant. As they both contained the same 48 basic-level categories, yet different exemplars of each, we were able to correlate the two context RDMs with each other to reduce visually-specific effects in the comparison (Fig. 5a, right panel). In order to compare the results to our within-exemplar reliability measures (Fig. 5b), we correlated the context RDMs across sessions (e.g. green-context RDM from session 1 with red-context RDM from session 2) and not within sessions. We hypothesized that, if there are fine-grained category representations in the patterns, they should show up as significant similarities between the two context RDMs.

Our results showed that there were indeed highly significant correlations between the two RDMs in lower and higher visual regions (EVC: mean r = 0.09 ± 0.008, LO: mean r = 0.07 ± 0.01, pFus: mean r = 0.09 ± 0.01, p < 0.001; Fig. 5b). We did not find any significant across-exemplar correlations in the representational geometries in HC or PRC (HC: mean r = 0.009 ± 0.006, p > 0.05; PRC: mean r = −0.002 ± 0.007, p > 0.05; Fig. 5b). In all ROIs, except for HC, the difference between the within-versus across-exemplar correlations were highly significant (p < 0.001).

To test whether reliability-based voxel selection introduced any specific bias, we repeated the Pearson correlation analysis on the beta arrays of voxels in a brainstem mask from *FSL* and selected the most reliable voxels (r >= 0.15). As expected, subsequent across-session RDM correlation analyses on these voxels indicated no significant effect (within-exemplar: mean r = 0.01 ± 0.007, p > 0.05; across-exemplar: mean r = −0.008 ± 0.006, p > 0.05), and no significant difference of using reliability-based voxel selection as compared to no selection (p > 0.05), demonstrating that no specific effect was introduced through this procedure.

These results suggest that the representational geometry of individual images is highly reliably represented in lower and higher visual regions, namely EVC and LOC, while across-exemplar representational similarities showed smaller yet significant effects. These results suggest that there is a stable representational geometry across different exemplars of the same basic-level category, consistent with our multivariate decoding results (Fig. 4). Post-hoc analyses on whole ROIs without reliability-based voxel selection revealed a similar pattern of results (Fig. 5c), yet with overall smaller effects, demonstrating no specific bias through the voxel selection strategy.

To assess the possibility that these across-exemplar effects are driven by shared low-level visual information, which we expected to be rather high within basic categories, we utilized variance partitioning to control for low-level visual feature effects. To this end, as a proxy of visual features, we took the RDM of the second convolutional layer of the DNN AlexNet layer that best explained the variance in EVC (mean r = 0.52; Cichy et al., 2016; Kubilius et al., 2016) and partialled out the explained variance. We refer to the correlations remaining after partialling out the early DNN layer as low-level-controlled (Fig. 5d), as the variance explained by the DNN layer is unlikely to capture mid- and high-level visual features (e.g. shape and texture) contained in the images. Please note that the term “low-level-controlled” does not imply full low-level control, which cannot be achieved without knowing the exact data generating process. These analyses were only applied to the ROIs showing significant across-exemplar correlations, namely EVC, LO and pFus (Fig. 5d). After partialling out the early layer RDM, the correlation values in EVC dropped significantly (mean difference = 0.03, p < 0.01; Fig. 5d). This indicates that the shared across-exemplar representations in EVC are partially dictated by low-level visual features. Looking further, in the LOC partialling out the variance explained by the early layer RDM did not lead to a significant decrease in the set of correlation values (mean difference < 0.01; p = 0.99; Fig. 5d). In line with the multivariate decoding results, this shows that there are some across-exemplar representations in LO and pFus that cannot be reduced to low-level visual features captured by the DNN layer but rather indicate a shared representation at a different level. To test whether these residual effects can be explained by shared mid-level visual features, we additionally partialled out the variance explained by a model of perceived shape similarity (Morgenstern et al., 2021). We found that after controlling for shape similarity between exemplars significant results remained significant and vice versa (low/mid-level-controlled; Fig. 5d). This shows that the across-exemplars representations in LO and pFus cannot be explained by the features of our models chosen to reflect low-level or mid-level visual information, suggesting that other, potentially more high-level information is encoded in these patterns.

### Similarities between neural and semantic knowledge model RDMs

The multivariate decoding as well as RSA results showed that the response patterns in LOC contain some form of object-specific information shared between different exemplars, with a stable representational geometry. While this is suggestive of semantic effects, this is only indirectly addressed with the tests of generalization between exemplars. To directly address the contribution of semantic knowledge to these patterns of brain responses, we related two numerical embeddings with fMRI response patterns using RSA (Fig. 6a), the THINGS behavioral similarity embedding (Hebart et al., 2020, 2023) and CSLB semantic feature norm (Devereux et al., 2014). Results are only reported for EVC, LO (Fig. 6b), and pFus, since HC and PRC showed no significant RDM generalization between exemplars (Fig. 5b).

**Figure 6:**
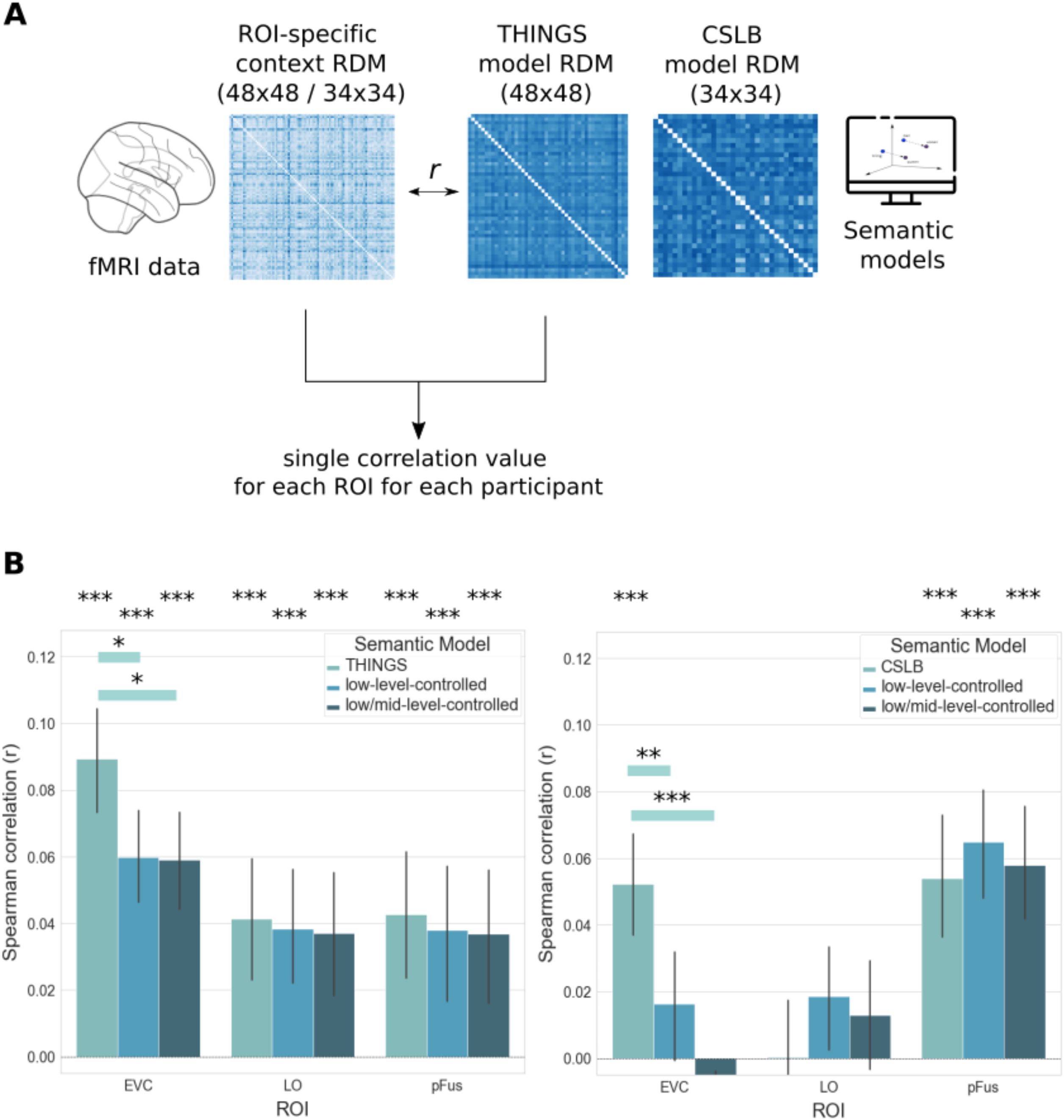
Comparison between neural voxel patterns and semantic knowledge models. **A)** Analysis approach for uncovering semantic information in the representational geometry of the fMRI signal. Two embedding model RDMs (THINGS embedding, CSLB) were correlated with the ROI-specific fMRI RDMs to detect significant correspondence. **B)** Spearman correlation values between the THINGS embedding (left) as well as the CSLB semantic (right) model RDMs and the ROI-specific pattern RDMs from each session and each participant. Effects are shown without variance partitioning (“THINGS”, “CSLB”), after solely partialling out the early convolutional layer RDM from AlexNet (“low-level-controlled”) as well as after additionally partialling out the shape model (“low/mid-level-controlled”). Results are shown only for ROIs with significant across exemplar reliability. Asterisks indicate significance when compared with zero at significance levels: *** p < 0.001, ** p < 0.01, * p < 0.05, Bonferroni-corrected. Error bars reflect 95% confidence intervals.

We found a highly significant correlation between the THINGS perceived similarity embedding RDM and the RDMs in EVC, LO and pFus (EVC: mean r = 0.09 ± 0.008, LO: mean r = 0.04 ± 0.009, pFus: mean r = 0.04 ± 0.009, p < 0.001; Fig. 6b, left). When partialling out the variance explained by the early layer DNN, which we again refer to as “low-level-controlled”, the correlation in EVC dropped significantly (mean difference in r = 0.03, p = 0.019) yet remained significant (mean r = 0.06 ± 0.007, p < 0.001; Fig. 6b) indicating that the semantic embedding partially explained variance dominated by low-level visual features in EVC. In LO and pFus, partialling out the DNN-related variance did not significantly change the correlations (p > 0.05; Fig. 6b), suggesting that the THINGS model captured information encoded in these regions that transcends the variance explained by the early DNN layer. In order to control for mid-level similarity effects, we next partialled out the variance explained by a model of perceived shape similarity (Morgenstern et al., 2021). We again found that after controlling for the variance explained by the shape similarity model the results of the embedding analysis remained significant (low/mid-level-controlled; Fig. 6b). These results complement our multivariate decoding and across-exemplar RSA results (Fig. 4, 5), suggesting that the information encoded in LO and pFus encompasses object-specific category information that extends beyond low-level and mid-level visual feature effects captured by our models.

Lastly, we tested to what extent the CSLB model based on semantic features could capture information encoded in the voxel patterns. The CSLB model significantly correlated with the patterns in EVC (mean r = 0.05 ± 0.008, p < 0.001; Fig. 6b, right), which again decreased significantly after partialling out the early DNN layer RDM (p < 0.01, low-level-controlled, Fig. 6b, right), leading to non-significant remaining effects (mean r = 0.02 ± 0.08, p = 0.25). The model was not able to explain any significant variance in LO (mean r = 0.00 ± 0.009, p = 0.99; Fig. 6b, right). However, in pFus the correlation with the CSLB model RDM was highly significant (mean r = 0.06 ± 0.009, p < 0.001) and was not significantly affected by partialling out the early DNN layer (p = 0.99) and remained significant (mean r = 0.06 ± 0.009, p < 0.001; Fig. 6b, right). Similar to the analyses above, partialling out the variance explained by our control model for mid-level shape similarity did not change the pattern of results and significant results remained significant (low/mid-level-controlled; Fig. 6c).

These findings further suggest that the response patterns in pFus encode higher-level category-specific information, in line with our previous decoding as well as RSA results.

### The effect of the color context on the representational geometry

Since our task contained a context component, it was important to rule out the possibility that the color context was adversely affecting the results. We tested for the influence of the context in two ways. First, we repeatedly mixed (n = 1000) the color contexts within each session across all stimuli and used this as a basis for forming new, mixed RDMs that were the same in content but mixed in color context. If the effects we reported were driven by color context, then shuffling the contexts should reduce or eliminate these results. However, the results remained unchanged after shuffling (p > 0.99), demonstrating that color context was not affecting the pattern similarity across sessions. Additionally, in order to test whether our analyses might miss obvious patterns in the similarity relation between stimuli, we conducted 2-dimensional metric multidimensional scaling (MDS) on the multivoxel patterns in all ROIs averaged across all participants and sessions. The results showed no obvious pattern related to color context (Fig. 7). To quantify these visually illustrated pairwise distances and test to what extent there were significant differences between the two populations, we computed a t-test comparing the within-with the across-context dissimilarity values within each participant. After controlling for multiple comparisons, we did not find significant differences between the dissimilarity values of within-versus across-context objects in any ROI (p > 0.05), again indicating that the context did not significantly influence the similarity values between the stimuli.

**Figure 7:**
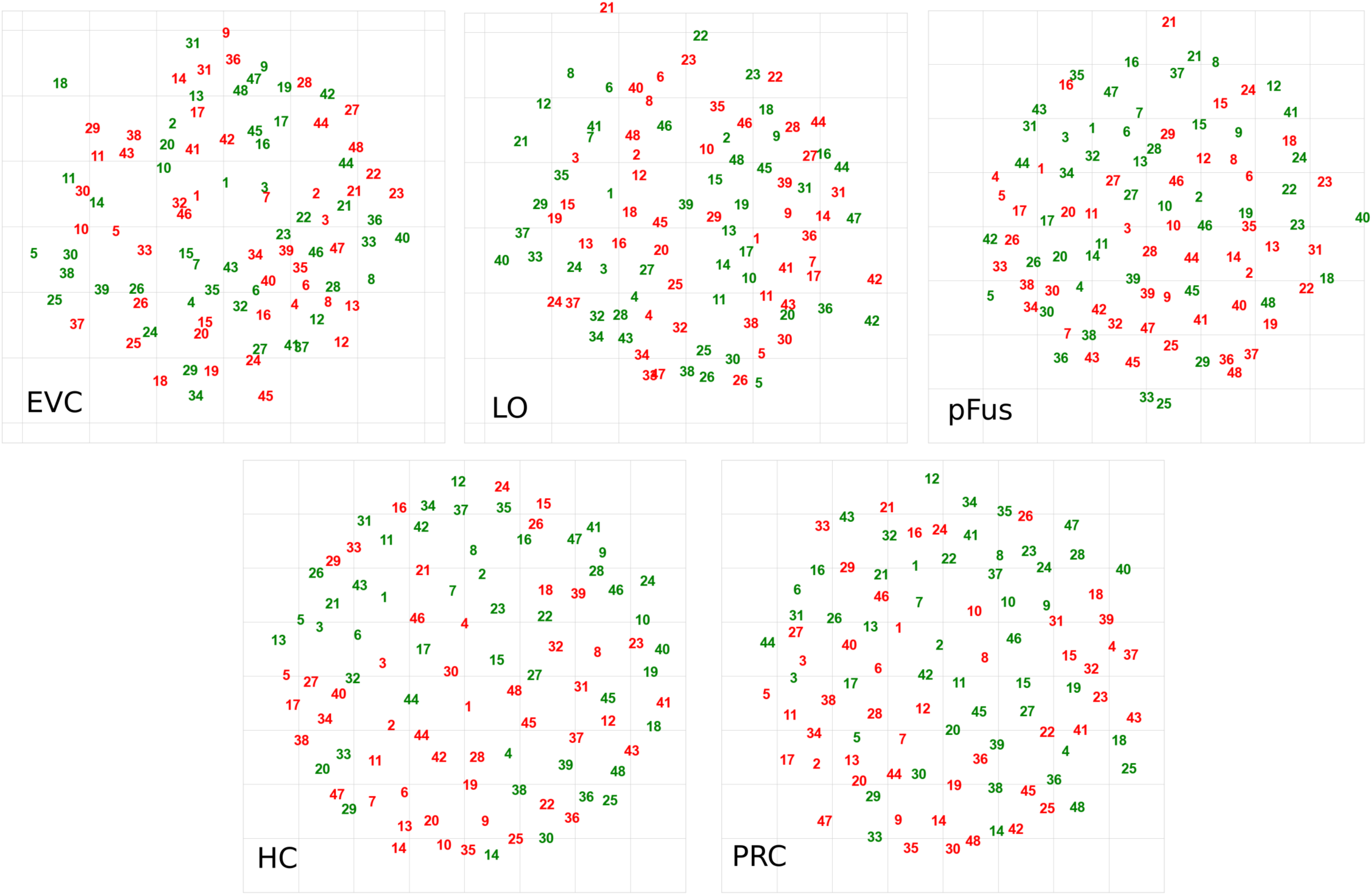
2D MDS representations on the multivoxel patterns. A 2-dimensional multidimensional scaling (MDS) display derived from the average RDMs across all participants and sessions for each ROI showing pairwise distances between all stimuli. The numbering ranges from 1 to 48 and displays the basic-level mammal category arranged in alphabetical order (as in Fig. 1a). The numbers are colored to indicate the color-context each stimulus belongs to.

Together, these results indicate that color context did not significantly affect the object-specific neural representations.

## Discussion

It is commonly assumed that fMRI studies can provide fine-grained information about basic-level objects in higher-level visual cortex (Eger et al., 2008; Cichy et al., 2011; Iordan et al., 2015). However, much previous work has left open to what degree previous findings could in part best be explained by coarse-grained category effects (e.g. animate vs. inanimate; mammal vs. insect) or by low-level visual features of individual objects (Andrews et al., 2010, 2015; Coggan et al., 2016; Bracci et al., 2017; Groen et al., 2017). In this context, the degree to which the global representational geometry remains stable or shifts between different object examples of the same basic-level category, has remained unclear. By putting together a novel stimulus set and leveraging different models, thereby carefully controlling for coarse categorical and visual feature effects, our study reveals three major findings. First, object-related information and the representational geometry along the ventral visual pathway are strongly dominated by image-specific effects. Second, there are much weaker yet reliable effects across images of the same object that cannot be explained by a low- or mid-level visual feature model. Third, a comparison of two embedding models of semantic knowledge showed that these remaining effects may, in part, be explained by high-level semantics.

Consistent with recent approaches of study designs in cognitive neuroscience (Naselaris et al., 2021; Hebart et al., 2020, 2023), our study emphasized extensive fMRI data sampling over multiple days to enhance statistical power and optimize the reliability of our findings. This is especially critical when looking for fine-grained representations encoded in the multivoxel patterns as inter-individual differences induced variability, and hence noise, which becomes particularly apparent at fine spatial scales (Weiner & Grill-Spector, 2012; Naselaris et al., 2021). Additionally, to maximize statistical power, we used strict behavioral inclusion criteria as well as refined model fitting (LS-S) and reliability-based voxel selection (Tarhan & Konkle, 2021).

Previous studies have supported the notion that visual features strongly drive responses along the ventral visual pathway, even in higher-level regions, making it challenging to disentangle visual from semantic information (Andrews et al., 2010, 2015; Baldassi et al., 2013; Long et al., 2017, 2018). In line with this, our decoding results revealed highly robust image-specific representations across scanning sessions. As expected, these effects were especially strong in early visual cortex (EVC) and slightly weaker in higher visual areas (LO and pFus, together LOC). Moreover, our RSA results showed a reliable image-specific representational geometry in EVC and, to a smaller extent, in LOC. Taken together, these results demonstrate that both the image-specific information and the image-specific representational geometry remained reliable across scanning sessions in early and later visual regions. Consequently, low-level visual feature effects appeared to be a strong and reliable driver of the representations in the regions.

Interestingly, our decoding results showed that reliable image-specific representations extended to regions in the anterior temporal lobe (HC and PRC), also mirrored in the reliable image-specific representational geometry. While in the past it has proven challenging to identify specific effects of visual stimuli in HC and PRC using more traditional vision tasks, it is possible that the color context task used in this study, in combination with the high within-participant statistical power, led to stronger engagement and more reliable effects of these regions, respectively. Beyond the potential for using this approach to studying visual representations in HC and PRC, these results reveal that low-level visual feature effects can in fact be reliably represented in such high-level brain regions and further confirm that supposedly high-level representations are significantly influenced by basic visual features (Bracci et al., 2017; Knapen, 2020; Silson et al., 2021). Thus, it is conceivable that effects previously related to higher semantic information (e.g. Clark & Tyler, 2014) are also affected by low-level visual information.

Category-specific representations should not be solely explainable by low-level visual features, but should be determined by distinct shared semantic features (Clarke & Tyler, 2015; Clarke, 2019). Prior fMRI research has suggested that there is category-specific information encoded in higher visual regions such as LOC (Eger et al. 2008; Cichy et al., 2011; Iordan et al., 2015). This information supposedly generalizes across exemplars of the same category, which suggests that these effects are best explained by shared visual features, semantics or a combination of both. Our results confirm and extend on these findings, in that we found small but significant shared representations between different exemplars of the same category in LOC. These effects are not significantly influenced by the visual similarity between exemplars, indicating that the representations indeed may contain category-specific information. Supporting these results, we found significant correlations between the fMRI response patterns in LOC and two embedding models that reflect semantic information (Devereux et al., 2014; Hebart et al., 2020, 2023). This indicates that the representations in LOC, beyond containing image-specific visual information, may carry some form of category-specific high-level information.

While our results clearly demonstrate that representations along the ventral visual stream are dominated by visual information, it remains possible that our control models did not capture some of the variance related to more low-level or mid-level visual features, and thus remaining across-exemplar information may be explained not by category-related but by visual information. In this context, a potential limitation of our study was that we did not select visually strongly different exemplars within-category, a methodological choice that could have further reduced the influence of visual features. Future models may show improved predictions of low-level and mid-level visual features and may thus help us more definitely tease apart the effects of low-level, mid-level and category-specific information in similar experimental designs. However, given the effectiveness of DNN models and semantic embeddings in capturing visual and semantic effects, respectively, it appears to be plausible that the remaining effects after low- and mid-level feature control are not primarily visual in nature but may also capture some degree of semantic effects.

The absence of high-level semantic information in the representations in PRC and HC could be explained by limitations of current fMRI techniques. First, the spatial and temporal resolution of fMRI might be too coarse to capture such fine-grained representations within such a homogenous object space. At the same time, image-specific information was captured to a certain degree, indicating that semantic information may be represented in a different format across voxels than visual information. Second, fMRI data from the anterior parts of the medial temporal lobe are known to contain weak and noisy signals (Weiskopf et al., 2006; Olman et al., 2009). Thus, such nuanced representations might not be picked up with fMRI in these regions even if the information is encoded there. An additional plausible explanation is that there is dynamic information flow between posterior and anterior regions of the ventral visual pathway, which in accordance allows for the dissociation between semantically similar objects (Clarke et al., 2011; Devereux & Clarke, 2018; Clarke, 2019). Capturing and discriminating between these dynamic representations requires data acquisition techniques with high temporal resolution. Future studies looking for basic-level object representations could use higher spatial resolution with fMRI (e.g. 7T) together with more dynamic neural recordings techniques (e.g. MEG), and could combine both techniques (Cichy et al., 201; Hebart et al., 2018).

In summary, our study went beyond previous work by using (1) a substantially larger group of categories with different exemplars, (2) a more fine-grained level of abstraction, (3) revealing not only generalizable decoding but also a stable representational pattern, and (4) demonstrating potential sources of this information using DNNs and semantic models (Eger et al. 2008; Cichy et al., 2011; Iordan et al., 2015). We deliberately chose categories of the class of animals, more specifically land mammals, as these stimuli have been extensively studied and we know them to be firmly represented along the ventral visual pathway (Kriegeskorte et al., 2008; Connolly et al., 2012, 2016; Sha et al., 2015; Bracci et al., 2019; Ritchie et al., 2021). Only including land mammals ensured that the effects between different exemplars of the category are indeed best explained by fine-grained semantics rather than coarse-grained categorical effects (e.g. having a face, degree of animacy; Sha et al., 2015; Proklova & Goodale, 2022). Our work highlights the dominance of image-specific visual feature representations yet demonstrates the presence of small, generalizable effects. This extends our understanding of the function of the ventral visual pathway and opens the door to a more detailed understanding of the nature of representations across these regions.

## Acknowledgement

We thank all the people who were so kind to read the manuscript and listen to project presentations, and whose constructive criticism and ideas have made this project possible.

